# Effects of Aerobic Exercise in Hepatic Lipid Droplet-Mitochondria interaction in Non-alcoholic Fatty Liver Disease

**DOI:** 10.1101/2023.01.31.526481

**Authors:** Juan Carlos Bórquez, Francisco Díaz-Castro, Francisco Pino-de La Fuente, Karla Espinoza, Ana María Figueroa, Inma Martínez-Ruíz, Vanessa Hernández, Iliana López-Soldado, Raúl Ventura, Alejandra Espinosa, Víctor Cortés, María Isabel Hernández-Alvarez, Rodrigo Troncoso

## Abstract

Lipid Droplets (LD) are highly dynamic storage organelles. In the liver, its accumulation causes non-alcoholic fatty liver (NAFL) that can progress to a more severe disease stage, nonalcoholic steatohepatitis (NASH). In hepatic and non-hepatic tissues LD interacts with mitochondria impacting lipid homeostasis. However, whether exercise modulates this interaction in the liver has not been studied yet. Our objective is to determine whether exercise modifies LD-mitochondria interaction in hepatocytes and if this interaction has an association with the severity of the disease. Two different models of NAFLD, a high fat diet (HFD) to evaluate NAFL and a methionine choline deficient diet (MCD) to evaluate NASH, were used to analyze the effects of aerobic exercise in the liver.

Our results in the NAFL model showed that exercise decreased the severity of the disease and improved physical capacity compared to sedentary HFD mice. In this regard, although exercise increased the number of LD in hepatocytes, LD were smaller in size than in the sedentary HFD mice. Notably, while sedentary HFD mice increased hepatic lipid droplet (LD)-mitochondria interaction, in exercised animals, this interaction was decreased. Additionally, exercise decreased the size of the LD bound to mitochondria, and this peridroplet mitochondria (PDM) exhibited higher basal respiration and ATP synthesis capacity than PDM from sedentary HFD mice. Besides, we found a positive correlation that predicts the severity of NAFL between LD-mitochondria interaction in the liver and plasmatic ALT transaminases. This correlation is also positive between hepatic LD-mitochondria interaction and the area under the glucose tolerance test curve in this model. Our results in the NASH model resemble, to a greater extent, what we observed in the NAFL model. In NASH, exercise also reduced collagen accumulation, decreased LD-mitochondria interaction, and reduced the size of LD coupled to mitochondria compared to sedentary MCD mice.

In all, our results show that aerobic exercise decreases LD-mitochondria interaction in hepatocytes and this interaction is associated with less severity of NAFL and NASH. We propose that exercise provokes an improvement of NAFLD by reduction of the hepatic LD-mitochondria interaction that in turn increase peridroplet mitochondria activity.

**Highlights:**

- Lipid droplet (LD)-mitochondria interaction is increased in high-fat diet-induced NAFLD and choline-methionine deficient diet induced-NASH.
- Exercise decreased LD-mitochondria interaction and is associated with reduced plasmatic ALT transaminase levels and glucose tolerance test in HFD induced-NAFLD.
- Exercise decreases LD-mitochondria interaction, decreasing peridroplet Mitochondria (PDM) with possible lipogenic function, which induces a decrease in the LD bound to mitochondria (M-LD) in HFD-induced NAFLD.
- Exercise decreased LD-mitochondria interaction and collagen accumulation in MCD induced-NASH.

## 1. Introduction

Non-alcoholic fatty liver disease (NAFLD) is the most common liver disease worldwide, affecting one-third of adults (1), with a high mortality risk due to its close association with obesity, insulin resistance, and cardiovascular disease (2). It is a clinical spectrum ranging from non-alcoholic fatty liver (steatosis, NAFL) and non-alcoholic steatohepatitis (NASH) to cirrhosis and hepatocellular carcinoma. NASH is distinguished from NAFL by the presence of hepatocyte injury (hepatocyte ballooning and cell death), inflammatory infiltrates, and/or collagen deposition (fibrosis) (3). Although several studies show that exercise attenuates the progression of NAFLD (4–8), the mechanism that explains this is not entirely clear. Some studies show that it is given that exercise decreases liver lipid accumulation (9,10). However, we and others had reported metabolic benefits without changes in hepatic lipid accumulation, body mass, or fat mass (11–14).

NAFLD is characterized by the excessive deposition of triglycerides (TG) as lipid droplets (LD) in the cytoplasm of hepatocytes (15). LD are not just lipid stores, they are now recognized as organelles that play a central role in lipid homeostasis by controlling the storage of fatty acids (FA) and their release from TG stores, thus preventing high cellular levels of toxic lipids. Structurally, they have a TG core enclosed by a phospholipid monolayer decorated by proteins (16). His life cycle includes LD biogenesis in the endoplasmic reticulum (ER), LD expansion by local synthesis of TG on the LD surface, and LD catabolism by lipolysis and lipophagy (15,16). Importantly, LDs have different features, including their size, number, surface proteins, and interaction with organelles, which are critical factors in determining the progression of the diseases (17).

Multispectral time-lapse imaging revealed that mitochondria are the second most common interaction partner for LD (18). In the liver, the role of lipid droplet-mitochondria interaction is unknown. As an approach, perilipin 5 (PLIN5) deletion in hepatocytes remodels lipid metabolism and causes hepatic insulin resistance in mice (19). By contrast, overexpression of PLIN5 in hepatocytes causes severe steatosis but, surprisingly, does not affect insulin sensitivity (20). In addition, we showed that oleic acid promotes more LD-mitochondria contacts than palmitic acid, reducing the charge over mitochondrial function in the hepatic cell line HepG2 (21). The LD-associated protein PLIN5 contains a single C-terminal domain not present in the other perilipins, which is necessary to establish contact between LD and mitochondria (22,23). On the other hand, the role of LD-mitochondria interaction may be tissue-specific. For example, brown adipose tissue LD-mitochondria interaction stimulates LD expansion. LD bound to mitochondria, or Peridroplet Mitochondria (PDM), have specific features that promote TG synthesis (24). By contrast, LD-mitochondria interaction in skeletal muscle is associated with using FA derived from LD catabolism. LD-mitochondria contact was greater in type 1 fibers (oxidative fiber) than in type 2 fibers of diabetic, nonobese diabetic, and lean subjects (25). In addition, exercise increased LD-mitochondria interaction and β-oxidation in the muscles of healthy subjects (26).

This study aimed to determine whether exercise modifies LD-mitochondria interaction in hepatocytes and its association with disease severity. Here, we showed that aerobic treadmill exercise decreased LD-mitochondria interaction in hepatocytes and is associated with reduced NAFL and NASH severity.

## 2. Material and Methods

### 2.1. Animals and diets

Animal care and procedures were approved by the Institutional Animal Care and Use Committee (CICUA) of the Universidad de Chile (20422–INT–UCH). Forty-five wild-type C57BL/6J male mice were housed in a temperature-controlled (20-23 º C) room with a 12:12-h light-dark cycle (6 AM - 6 PM) and *ad-libitum* access to chow food and tap water.

Cohort 1. At 4 weeks of age, C57BL/6J mice were randomly assigned to a control diet (CD. 20% protein, 70% carbohydrate,10% fat in total calories D12450J, Research Diets, New Brunswick, NJ, United States) or high-fat diet (HFD. 5.21 kcal/g. 20% protein, 20% carbohydrate, and 60% fat in total calories. D12492, Research Diets, New Brunswick, NJ, United States) for 8 weeks. Then, mice were subdivided into 4 groups: CD, CD + exercise, HFD, and HFD + exercise, and followed for additional 4 weeks.

Cohort 2. Ten C57BL/6J male mice (8 weeks of age) were randomly assigned to CD or a methionine choline-deficient diet combined with a high-fat diet (MCD, A06071305 MCD with 45% HFD from Research Diets) and supplemented with 0.1% L-methionine on drinking water for 3 weeks.

Cohort 3. Fourteen C57BL/6J male mice (8 weeks of age) were fed with MCD and randomly assigned to exercise or kept sedentary for 4 weeks.

All mice were sacrificed after 6 hours of fasting and euthanized anesthesia with isoflurane at 5% (zoetis, Parsippany, NJ, United States). Sample blood was rapidly drawn from the abdominal vena cava into pre-cooled syringes treated with ethylenediaminetetraacetic acid (EDTA) and stored on ice. Blood was centrifuged (3500 rpm for 10 min, 4 °C), and plasma was removed and frozen (−80 °C) until analysis.

### 2.2. Treadmill exhaustion test and exercise training protocol

All mice were acclimatized on the treadmill for 3 days. On the first and second days, mice ran for 10 min at a speed of 0.5 km/h (14 cm/s) with a slope of 0°. On the third day, the mice ran for 10 min at 0.5 km/h (14 cm/s), 5 min at 0.6 km/h (17 cm/s), and 5 min at 0.8 km/h (22 cm/s) with a 0° slope. After 2 days of rest, the mice performed a graded exercise test until voluntary exhaustion. The test began at 0.5 km/h (14 cm/s) with a slope of 0°for 5 min, followed by a speed increase of 0.1 km/h (3 cm/s) every 3 min until the animal could no longer maintain treadmill speed. The same protocol was performed at the end of the period of training.

Exercise training on the treadmill (LE8710; Harvard Apparatus) was performed for 4 weeks, 5 times per week for 1 hour each session at 60–65% of the maximal speed reached on the treadmill exhaustion test. Finally, mice were sacrificed at least 72 h after the last exercise session.

### 2.3. Grip Strength Test

Total strength was determined by an all-grip strength test measured in a force gauge (Force Gauge, China). Mice were placed horizontally over the metal grid and pulled by the tail continuously until they could not hold on anymore. This procedure was repeated three times for each mouse to obtain a mean value. Strength determination was done before and after the training exercise. Grip strength was expressed in N (Newtons) and normalized by body weight (N/g).

### 2.4. Glucosa tolerance test

An intraperitoneal glucose test (GTT) was performed at weeks 8 and 12 of treatment. Mice were fasted for a minimum of 6 h, with free access to water. They were given an intraperitoneal injection of glucose (2g/kg body weight) dissolved in saline. Blood glucose levels were measured from the tail vein at 0, 10, 20, 30, 60, 90, and 120 min using an Accu-Check Performa glucometer (Roche Diagnostic, Mannheim, Germany).

### 2.5. Biochemical analysis

Plasma concentrations of GOT/AST (U/L), total cholesterol(mg/dL), c-HDL (mg/dL), and triglycerides (mg/dL) were measured using a Spotchem II Kenshin-2 kit (77188, Arkray, Kyoto, Japan) in a SpotChem Analyzer (Arkray), according to the manufacturer*’*s instructions. Plasma insulin levels were determined by an enzyme-linked immunosorbent assay (ELISA) kit (10-1247-01, Mercodia, Uppsala, Sweden).

### 2.6. Histology analysis

Liver sections were stained with hematoxylin-eosin and Sirus Red. Images were visualized under a light microscope (CX22, Olympus, Tokyo, Japan) and taken 20x fields from the same liver lobe of each animal using the Toup View software (ToupTek Photonics, Zhejiang, China). The number and area of lipid droplets were measured with Image J program version 1.51 (National Institutes of Health, Bethesda, MD, USA) and quantification of Sirus red with QuPath software (QuPath developers, University of Edinburg)(27).

### 2.7. Transmission Electron microscopy

Liver samples were processed as previously described (28). Briefly, liver samples were sectioned in small fragments with a razor blade to 1 mm and then fixed in 2.5% glutaraldehyde 2% paraformaldehyde solution 0.1 M at 4 C for 2 h. The samples were washed three times with 0.1 M phosphate buffer. Following post-fixation in 1% osmium tetroxide in 0.1 M phosphate buffer at 4º C for 2 h, they were washed with highly pure water and kept overnight in 0.1Mphosphate buffer. Afterward, samples were dehydrated at 4º C under shaking in graded acetone solutions (50%, 70% and 90%) in highly pure water. They were then gradually infiltrated with Eponate 12 Resin (TED PELLA 18010), and polymerization of the resin was processed for 48 h at 60ºC. Thin sections (50 nm) were cut using a Leica EM UC6 (Leica, Vienna) and mounted on bare 200-mesh copper grids. Tissue sections were stained with uranyl acetate 2% for 30 min, washed with highly pure water, incubated for 5 min with lead citrate, and air-dried. Sample sections were examined on a transmission electron microscope (Tecnai T12 at 80 kV; Philips. Microscopy Facility Universidad Católica; Chile) and transmission electron microscope (Jeol EM J1010 100 kV, CCIT of Universitat de Barcelona). For HFD experiments, 112 electron micrographs were obtained at 8,200 magnification in a randomized, systematic order, and 135 electron micrographs were obtained at 12000x magnification for MCD experiments.

The identification of mitochondria on micrographs was based on visualizing a spherical or elongated shape in the cytosol, double membrane, and higher electron density (29). Identification of LDs was based on visualizing LDs that are grayish-white in appearance, spherical with a diffuse rim (absence of clearly visible membrane), and LDs with irregular membranes as previously described (25).

Analyses of the liver were performed using Fiji/ImageJ software (National Institutes of Health, Bethesda, MD). Since the LD-mitochondria interaction is described as adherence rather than proximity (25,30), physical contact between the edge of the LD surface and the outer membrane of the mitochondrion by electron microscopy was considered LD-mitochondria interaction. In addition, a high electron density is often observed at the site of contact between the two organelles. It is presented as the absolute number of contacts (number of contacts per field), number of contacts relative to the total number of LD, and number of contacts relative to the total number of mitochondria. Mitochondria and LD size were calculated as the mean area of each micrograph. Perimeter, diameter, aspect ratio (length-to-width ratio), and the number of mitochondria were calculated for each micrograph using the Fiji/ImageJ software (particle analysis). Mitochondrial density was calculated as the sum of the area of each mitochondrion divided by the total area of the electron micrograph.

### 2.8. Isolation Lipid Droplet

Hepatic LD purification were isolated as previously described (31). Briefly, after liver perfusion with 0.9% NaCl and 0.1% EDTA solution, the liver was placed on a Petri dish, chopped with a scalpel for two min, and transferred into a Dounce tissue grinder at a ratio of 1 g of tissue to 3 ml of homogenization buffer (25mMTris-HCl, pH 7.5, 100mMKCl, 1 mM EDTA, and 5 mM EGTA). After 6 up-and-down strokes a 200 rpm, the liver homogenate was centrifuged at 500g for 10 min at 4°C. 2.5 ml of the resulting post-nuclei supernatant (PNS) was mixed with an equal volume of 2.5 M sucrose and placed at the bottom of a sucrose step gradient of 25%, 15%, 10%, and 5% (w/v) sucrose in homogenization buffer, with an additional top layer of 25mMTris-HCl, pH 7.5, 1 mM EDTA and 5 mM EGTA, and centrifuged at 12,000g for 1 hour at 4°C (SW-41Ti rotor, Beckman Coulter, Pasadena, California). Six or seven fractions were collected from the top. To purify LDs, the LD fraction on the top of the gradient was recovered and concentrated by re-floating LDs at 16,000g for 10 min at 4°C. The lower phase containing the excess buffer was removed by aspiration with a syringe, and four volumes of ice-cold acetone were added to precipitate proteins and kept 48 hours at -20°C. The samples were centrifuged at 16,000g for 10 min at 4°C, the pellet was washed with cold acetone 3 times, air-dried, and reconstituted with 10 mM Tris-HCl, pH 7.5. After sonication, protein concentration was quantified by Novagen BSA assay (71285-3, Sigma-Aldrich).

### 2.9. Isolation mitochondria subpopulation

Cytosol mitochondria (CM) and peridroplet mitochondria (PDM) were isolated as previously (24,30,32). Briefly, After ∼1.5 g liver was washed with PBS, the liver was placed on a petri-dish, chopped with a scalpel, and transferred into a Dounce tissue grinder and added 7 mL de mitochondrial isolation buffer + BSA (50 mM sucrose, 5 mM HEPES, 2 mM EGTA, 2 mg/200mL fatty acid-free BSA). After 10 up-down strokes at 200 rpm in a homogenizer (RZR 2020, Heidolph), the liver homogenate (aliquot of 200 uL was collected) was centrifuged at 900 g for 11 min at 4°C. The resulting lipid and aqueous phases were separated, and centrifugation was repeated. The lipid and aqueous phases were recovered from each tube (200 uL of each phase were collected), and both were centrifuged at 9000xg for 18 min at 4º C. (type 90 Ti fixed-angle rotor. Centrifuge L-90K ultracentrifuge). After recovering the lipid phase (stripping fat phase, 200 uL aliquot was collected) from one tube and the aqueous phase (cytosol, 200 uL aliquot was collected) from the other tube, the resulting pellets from both tubes (PDM and CM) were resuspended with mitochondrial isolation buffer without BSA. Aliquots saved in the process were analyzed by immunoblot to determine the purity of the isolation of mitochondrial subpopulations.

### 2.10. Oxygen Consumption Assessment in isolate mitochondria subpopulation

CM and PDM were resuspended in Mitobuffer (50 mM sucrose, 5 mM HEPES, 2 mM EGTA). The suspension was placed in a chamber coupled to Clark*’*s electrode (Oxygraph, Hansatech Instruments, Norfolk, UK). Oxygen levels were recorded for 3 min intervals at baseline (5 mM glutamate/2.5 mM malate), maximal capacity synthesis ATP (150 μM, ADP) and non-ATP-associated (1 μM Oligomycin, 75351, Sigma-Aldrich) conditions. Oxygen consumption rates were calculated and normalized for protein content by Novagen BSA assay (71285-3, Sigma-Aldrich). Results are shown as fold change with respect to CM baseline condition.

### 1.1. Western Blot

For whole liver lysate analyses, it was lysed with T-PER lysis buffer (78510, Thermo Fisher Scientific) supplemented with commercial phosphatases and protease inhibitors (04906845001 and 11836153001, Roche, Basel, Switzerland) and centrifuged at 1,000 g for 10 min at 4◦C. The supernatant protein content, LD fraction, and hepatic mitochondrial subpopulation were assessed using the Novagen BCA assay and denatured in an SDS buffer. 30-50 micrograms of proteins were separated by SDS– PAGE (10% polyacrylamide), electrotransferred onto PVDF membranes, and blocked with 5% nonfat milk in 0.1% v/v Tween 20 Tris-buffered saline (pH 7.6). Membranes were incubated in 0.1% v/v Tween 20 Tris-buffered saline with 1:1000 primary antibodies against PLIN5 (1:1000,26951-1-AP, Proteintech), mtHsp70 (1:1000, #MA3-028, Thermo FisherScientific), ATGL (sc-365278, Santa Cruz Biotechnologies), OXPHOS (1:1,000, ab110413, Abcam, Cambridge, United Kingdom) at 4◦C overnight and re-blotted with horseradish peroxidase-linked secondary antibody (1:1000). Finally, the bioluminescent signals were detected using ECL-plus reagent (DW1029, Biological Industries) and C-DiGit Blot Scanner (LI-COR, Lincoln, NE). Bands were analyzed using Fiji/ImageJ software (National Institutes of Health, Bethesda, MD) and quantified using scanning densitometry.

### 1.2. Statistical Analysis

A two-way analysis of variance (ANOVA) was used for statistical analysis, followed by a Tukey post-hoc test for multiple comparisons. Also, the Pearson correlation coefficient was performed, and p< 0.05 were considered significant. Results were expressed as mean ± standard error of the mean (SEM). The statistical analysis was performed using GraphPad Prism 7 software.

## 3. Results

### 3.1. Exercise decreased severity and improved physical capacity in HFD-induced NAFL mice model

To explore the role of aerobic exercise in HFD-induced NAFL, wild-type mice were fed with a control diet (CD) or high fat diet (HFD) during 8 weeks. Then mice were randomized to keep sedentary or aerobic exercise (CD+Ex, HFD+Ex) during 4 weeks more (**Supplementary fig. 1**). As expected HFD mice increased body weight (**Fig. 1A-B**), and this was attributable to increased caloric intake (**Fig. 1C**). In those conditions, HFD mice exhibit increased glucose intolerance (**Fig. 1F-G**) and insulin resistance (**Fig. 1H-J**), altered serum lipoprotein levels (**Fig. 1K-N**), higher liver transaminase levels (**Fig. 10-P**) and higher epididymal white adipose tissue weight (eWAT) (**Supplementary fig. 2D-F**) compared to CD. In addition, HFD mice decreased their physical capacity, given by lower maximal speed in the treadmill test (**Fig. 1E**) and lower grip strength (**Fig. 1D**). By contrast, exercise decreased NAFL severity and improved physical capacity in NAFLD. Although HFD and HFD+Ex mice did not change their body weight gain during exercise or caloric intake (**Fig. 1B-C**), exercise reversed and attenuated the HFD-induced decrease in maximal speed and strength, respectively (**Fig. 1E-D**). Moreover, HFD+Ex mice increased muscle fiber size (**Supplementary fig. 2C**), accounting for an improvement in muscle function. In addition, exercised mice decreased glucose intolerance and insulin resistance (**Fig. 1F-J**). They also show improvements in serum lipoprotein levels, decreasing total cholesterol levels (**Fig. 1K**), and serum triglycerides (**Fig. 1N**). In addition, exercise decreased liver damage by diminishing the levels of hepatic serum transaminases (**Fig. 10-P**) compared to HFD mice. Despite no changes in eWAT (**Supplementary fig. 2D-E**), exercise reduced adipocyte size compared to HFD (**Supplementary fig. 2D-F**).

**Figure 1.**
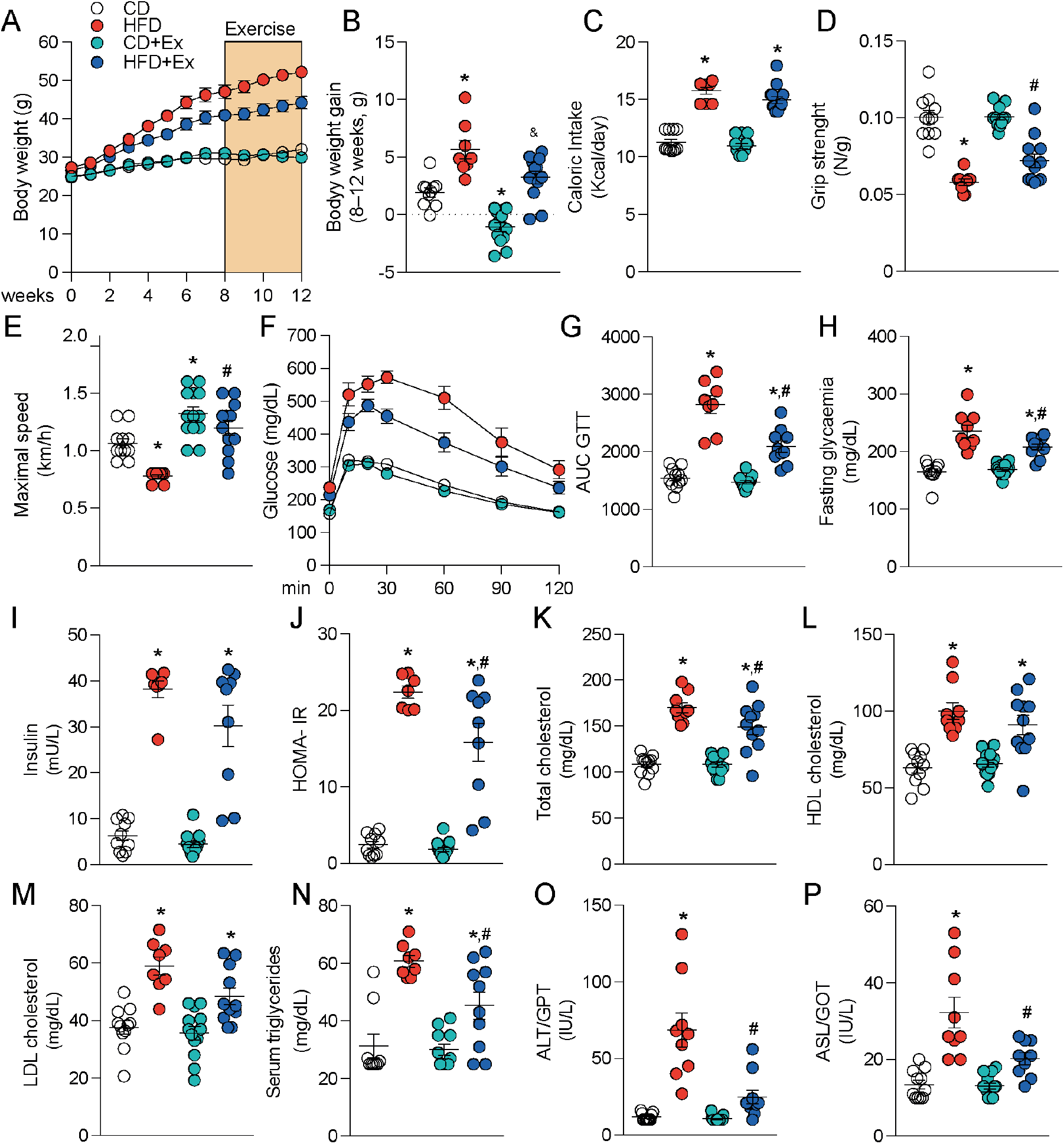
Exercise decreased the severity of HFD-induced NAFL. **A)** Body weight. **B)** Body weight gain. **C)** Caloric Intake. **D)** Grip strength. **E)** Maximal speed. **F)** Glucose tolerance test (GTT) **G)** Area under the curve (AUC) of GTT. **H)** Fasting glycemia. **I)** Serum insulin. **J)** Homeostatic Model Assessment of Insulin Resistance (HOMA-IR). **K)** Total cholesterol. **L)** HDL cholesterol. **M)** LDL cholesterol. **N)** Serum triglycerides. **O)** Alanine aminotransferase. **P)** Aspartate transaminase. Data presented as mean SEM. n=9-13 mice per group. 2-way ANOVA followed by Tukey*’*s post-test. Statistical significance p=<0.05; **\***vs CD, ^**&**^vs CD+Ex, ^**#**^vs HFD. CD: control diet; CD+Ex: control diet + exercise; HFD: high-fat diet; HFD+Ex: high-fat diet + exercise.

### 3.2. Exercise reduces hepatic LD size without impact in hepatic mitochondrial morphology in HFD-induced NAFL mice model

Next, we determine the effect of exercise on the liver. HFD+Ex mice showed a reduction in weight (**Fig. 2B**) and a microvesicular pattern in the liver (**Fig. 2C**) compared to sedentary HFD. Large LDs are a hallmark of NAFLD, associated with worse severity of the disease (33). However, there were no changes induced by exercise in the NAFLD score (**Fig. 2D**). In addition, HFD+Ex mice showed a greater number of LDs in hepatocytes (**Fig. 2E**), but they were smaller in size than in HFD mice (**Fig. 2F**), without changes in the protein levels of PLIN5 (**Fig. 2H**), which is an integral protein of the LD that allow the contact with mitochondria (34).

**Figure 2.**
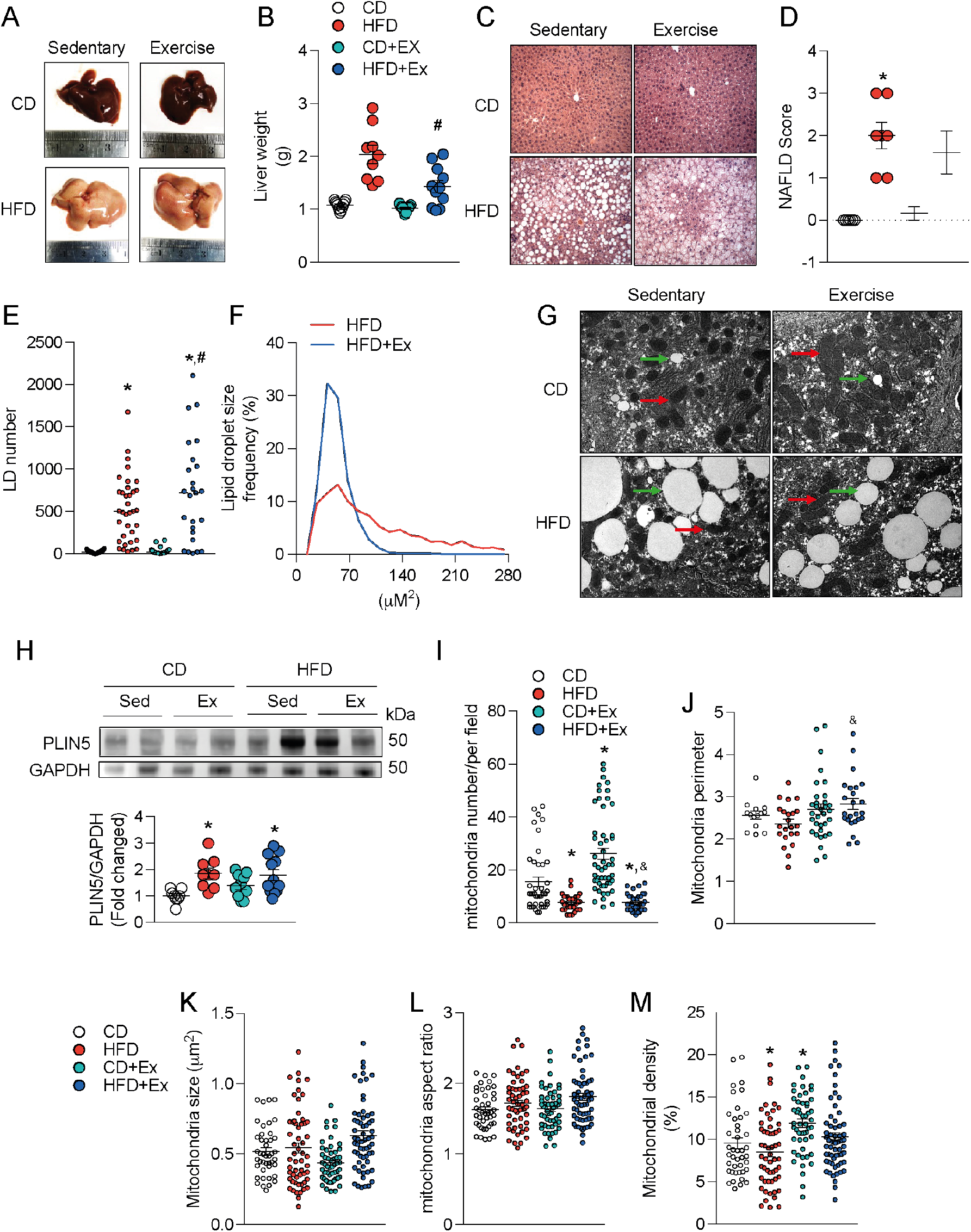
Exercise increased the frequency of small LD in HFD induced-NAFLD. **A)** Representative images of mice liver. **B)** Liver weight. **C)** Liver (H&E stain histology, 20x). **D)** NAFLD score. **E)** Lipid droplet number in the liver. **F)** Lipid droplet size frequency in the liver. **G)** Representative transmission electron microscopy (TEM) micrograph of mice liver (8200x, bar scale 2 μm). **H)** Perilipin 5 levels (PLIN5) by western blot **I)** Mitochondria total number. **J)** Mitochondria total perimeter. **K)** Mitochondria total size. **L)** Mitochondrial total aspect ratio. **M)** Mitochondrial total density from electron microscopy analyses (n=4-5 by group, each point means one micrograph analyzed). Data presented as mean SEM. n=9-13 mice per group.). 2-way ANOVA followed by Tukey*’*s post-test. Statistical significance p=<0.05; **\***vs CD, ^**&**^vs CD+Ex, ^**#**^vs HFD. CD: control diet; CD+Ex: control diet + exercise; HFD: high-fat diet; HFD+Ex: high-fat diet + exercise.

Compromised mitochondria in hepatocytes are also linked to worse NAFLD severity (35). In this regard, the number and density of total mitochondria decreased in HFD compared to CD (**Fig. 2I-M**). However, HFD+Ex showed no change in the number, perimeter, size, elongated morphology, and density of mitochondria compared with HFD mice (**Fig. 2I-M**). In contrast, the number and density of mitochondria increased in CD+Ex compared to CD (**Fig. 2I-M**). Therefore, exercise did not modify the number, density, or morphology of total mitochondria in NAFLD.

### 3.3. Exercise decreases Lipid Droplet-Mitochondria interaction in hepatocytes and is associated with reduced NAFL severity

To understand how exercise improves NAFL, we focused on hepatic LD-mitochondria interaction as a possible mechanism. Indeed, our results confirm that HFD increased LD-mitochondria interaction compared to CD in liver. Meanwhile, HFD+Ex decreased this interaction compared to HFD (**Fig. 3A-D**). In addition, a positive correlation was observed between LD-mitochondria interaction and ALT transaminase (**Fig. 3E**) and AUC values of GTT (**Fig. 3F**). These results suggest that exercise decreases LD-mitochondria interaction in hepatocytes, correlating with reduced severity of NAFL. Unexpectedly, in CD+Ex increased LD-mitochondria interaction compared to CD was observed, opposite to HFD mice (**Fig. 3A-D**).

**Figure 3.**
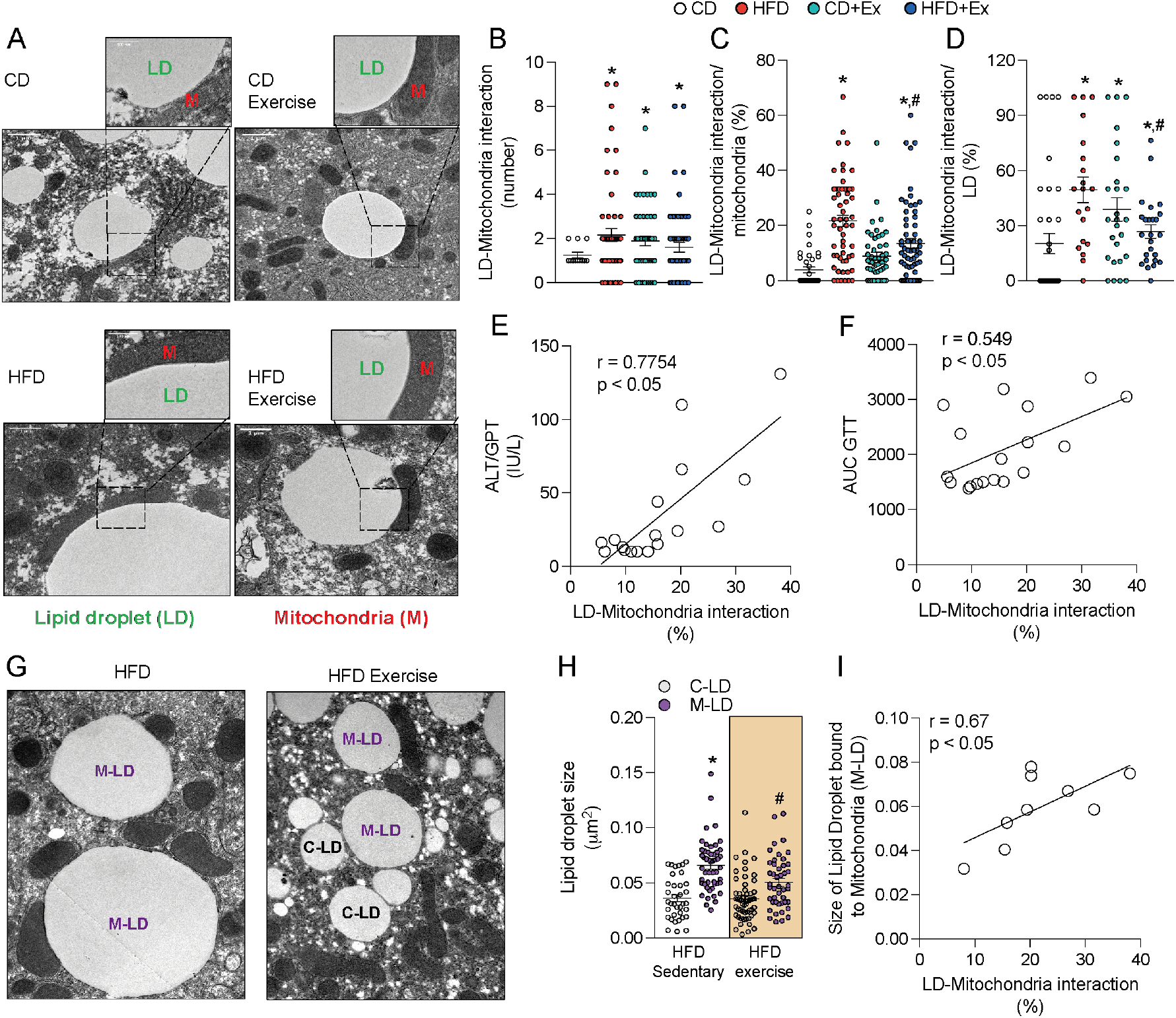
Exercise decreased Lipid droplet-mitochondria interaction in hepatocytes it is associated with reduced ALT transaminase levels and area under the curve of glucose tolerance test in HFD induced-NAFLD. **A)** Representative TEM micrograph of mice liver (big micrograph; 8200x, bar scale 1 μm. small micrograph; 28000x, bar scale 500 nm). **B)** Number of Lipid droplet (LD)-mitochondria interaction. **C)** LD-Mitochondria interaction by the total number of mitochondria. **D)** LD-Mitochondria interaction by the total number of LD. Correlation between LD-Mitochondria interaction/ total number of mitochondria (%). **E)** ALT transaminase. **F)** Area under the curve (AUC) of glucose tolerance test (GTT). **G)** Representative TEM micrograph of mice liver (big micrograph; 8200x). **H)** LD size. **I)** Correlation between size of LD bound to mitochondria (M-LD) and LD-Mitochondria interaction/total number of mitochondria (%). Data presented as mean SEM. n=4-5 by group, each point means one micrograph analyzed by TEM. In correlation graphs, each point means one mouse. 2-way ANOVA followed by Tukey*’*s post-test and Pearson correlation coefficient were performed. Statistical significance p=<0.05; **\***vs CD, ^**&**^vs CD, ^**#**^vs HFD. CD: control diet; CD+Ex: control diet + exercise; HFD: high-fat diet; HFD+Ex: high-fat diet + exercise; M-LD: Lipid droplets bound to mitochondria; C-LD: Lipid droplet non-bound to mitochondria.

Next, the role of LD-mitochondria interaction was explored to explain its positive correlation with NAFL severity. For this purpose, LD bound to mitochondria (M-LD) were studied. HFD increased M-LD size compared to cytosolic or non-mitochondria-bound LDs (C-LD) (**Fig. 3H**). Furthermore, a positive correlation was observed between the size of M-LD and LD-mitochondria interaction (**Fig. 3I**). In addition, in HFD+Ex mice the size of LD-M was decreased (**Fig. 3H**). These results suggest that LD-mitochondria interaction promotes LD expansion. Interestingly, exercise did not change the size of C-LD. Thus, exercise exclusively decreases LD-M size in NAFL.

To gain insight into the features of mitochondria in contact with LD, we studied the morphology and function of peridroplet mitochondria (PDM) and cytosolic mitochondria (CM). In electronic microscope images, PDM were found in lower proportion relative to CM in both HFD and HFD+Ex conditions (**Fig. 4A-B**). Also, PDMs show more elongated morphology compared to CM (**Fig. 4C**). In HFD+Ex, increased elongated morphology was observed in PDM relative to CM (**Fig. 4C**), suggesting that exercise increases the elongated morphology of PDMs in NAFL. Then we decided to isolated PDMs and CMs from liver tissue to further explore this effect. Despite not observing changes in protein levels of mitochondrial complexes in PDM from HFD and HFD+Ex (**Fig. 4D-I**), PDM from the HFD+Ex group exhibited higher basal respiration and ATP synthesis capacity than PDM from HFD (**Fig. 4J**). Therefore, exercise increases the ATP synthesis capacity of PDM in NAFL.

**Figure 4.**
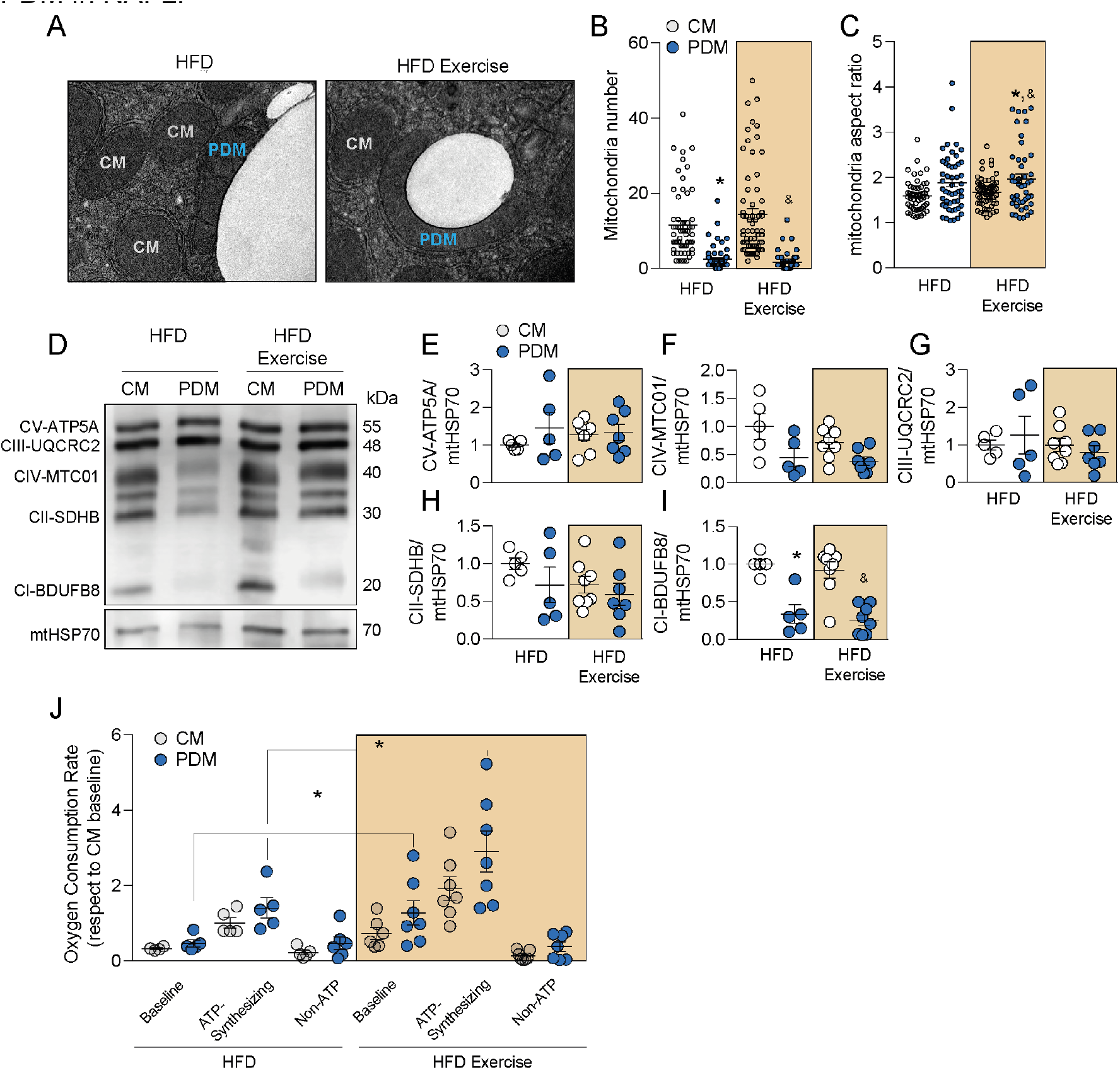
Exercise decreased size of LD bound to mitochondria and increased the ATP synthesis capacity of peridroplet mitochondria in HFD induced-NAFLD. **A)** Representative TEM micrograph of mice liver (12000x, bar scale 2 μm). **B)** Mitochondria number. **C)** Mitochondria aspect ratio. **D)** Representative images of western blot. **E)** Mitochondrial complex V. **F)** Mitochondrial complex IV. **G)** Mitochondrial complex III. **H)** Mitochondrial complex II. **I)** Mitochondrial complex I. **J)** Oxygen consumption under 5 mM glutamate/malate 2.5 mM (baseline condition), 300 μM adenosine diphosphate-ADP (maximum capacity to synthesize ATP), and 1 μM Oligomycin (non-ATP-associated). Data presented as mean SEM. n=4-5 by group (TEM analyses). n=5-7 by group (western blot and respirometry experiments). 2-way ANOVA followed by Tukey*’*s post-test. Statistical significance p=<0.05; **\***vs CD, ^**&**^vs CD. CM: cytosolic mitochondria; PDM: peridroplet mitochondria.

### 3.4. Hepatic LD-mitochondria interaction was increased in MCD induced-NASH mice model

Next, we were interested in observing the capacities of exercise in a harsh model of the disease. For that, the role of LD-mitochondria interaction in NASH was tested. For this purpose, mice were fed a methionine-choline deficient diet (MCD) supplemented with 0.1% of methionine in the drinking water to avoid weight loss (28) for 3 weeks to develop NASH, using as control, mice fed with CD (**supplementary fig. 3**). As expected, the MCD group did not change their body weight (**Fig. 5A**), but increased LD accumulation and collagen deposition in the liver (**Fig. 5B-E**) compared to CD. The MCD group increased LD-mitochondria interaction compared to CD (**Fig. 5F-I**). In addition, the MCD group showed higher levels of PLIN5 in the whole liver lysate (**Fig. 5J**) and isolated LD (**Fig. 5K**). Therefore, LD-mitochondria interaction also increases in the NASH model.

**Figure 5.**
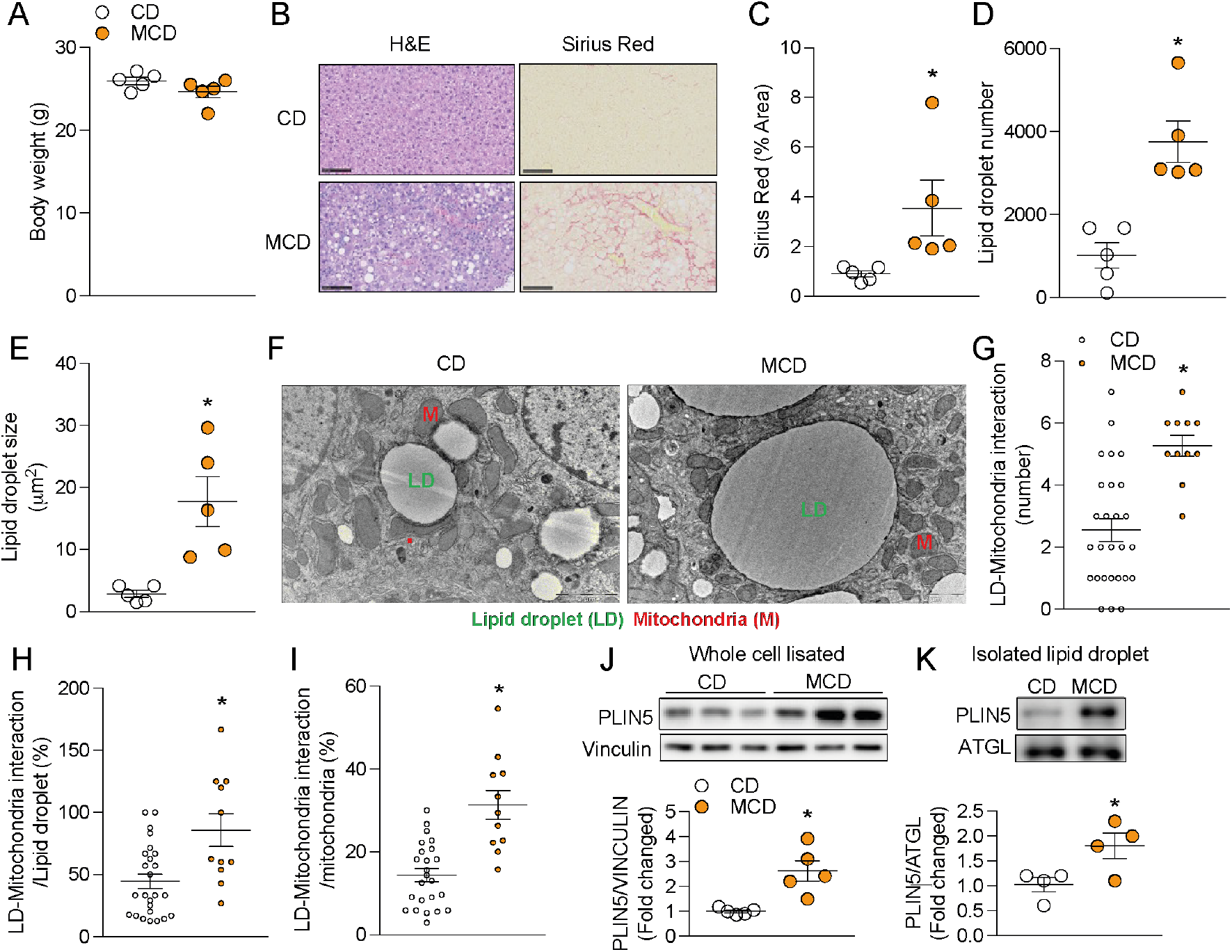
LD-mitochondria interaction was increased in MCD induced-NASH. **A)** Body weight. **B)** Representative images of mice liver (H&E and SR stain histology, 20x, bar scale 100 μm). **C)** Sirus Red. **D)** Lipid droplet number. **E)** Lipid droplet size. **F)** Representative TEM micrograph of mice liver (12000x, bar scale 2 μm). **G)** Number of Lipid droplet (LD)-mitochondria interaction. **H)** LD-Mitochondria interaction by the total number of LD. **I)** LD-Mitochondria interaction by the total number of mitochondria. **J)** PLIN5 levels in whole cell lysates by western blot. **K)** PLIN5 levels in isolated lipid droplets. Data presented as mean SEM. n=5 by group. In G), H), and I), each point means one micrograph analyzed by TEM. A t-student test was performed. Statistical significance p=<0.05 *v/s CD. CD: Control diet; MCD: methionine-choline deficient diet.

### 3.5. Exercise decreased hepatic LD-mitochondria interaction and liver collagen deposition in MCD induced-NASH mice model

Next, the effect of exercise on the severity of NASH was evaluated. For this purpose, a new cohort of mice were fed with MCD for 4 weeks and randomly designated as sedentary (MCD) or exercised (MCD+Ex) since the beginning of the diet treatment (**Supplementary fig. 4**). In this model, exercise reduced body weight gain (**Fig. 6A**), without differences in food intake (**Fig. 6B**), maximal speed (**Fig. 6C**) and intrahepatic triglycerides (**Fig. 6D**) compared to MCD. However, exercise reduces collagen accumulation (**Fig. 6E-F**). In addition, the MCD+Ex group has a higher number and reduced size of LD (**Fig. 6E, G-H**), with no changes in PLIN5 protein levels (**Fig. 6I-J**). Likewise, the MCD+Ex group shows a higher number and density of total mitochondria but of smaller perimeter and size (**Fig. 6K-P**). Taken together, exercise decreases the severity of NASH.

**Figure 6.**
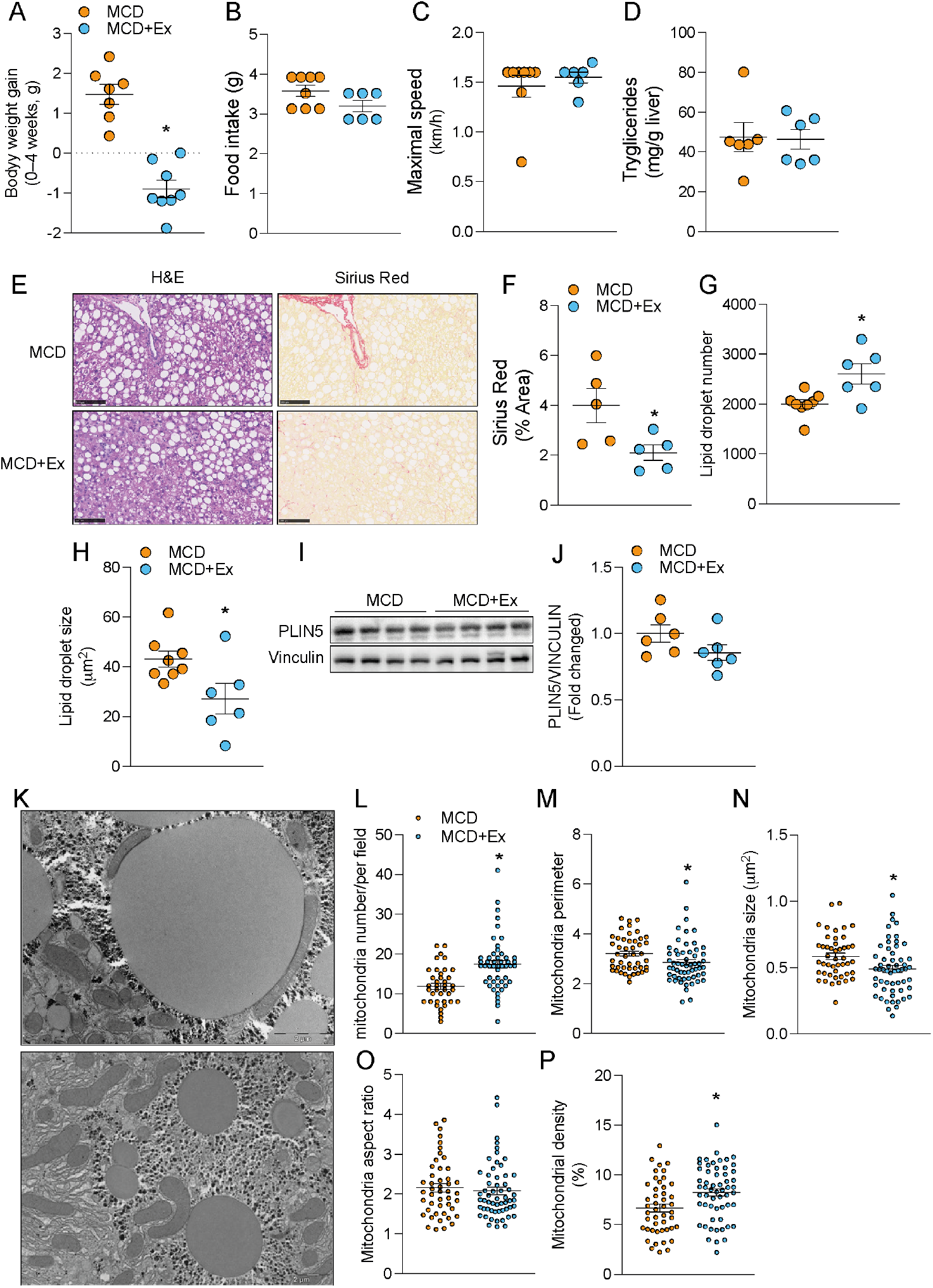
Exercise reduced severity and collagen accumulation in MCD induced-NASH. **A)** Body weight gain. **B)** Food intake. **C)** Maximal speed. **D)** Serum triglycerides. **E)** Representative images of mice liver (H&E and SR stain histology, 20x, bar scale 100 μm). **F)** Sirus Red. **G)** Lipid droplet number **H)** Lipid droplet size. **I), J)** and **K)** PLIN5 levels by western blot. **L)** Representative TEM micrograph of mice liver (12000x, bar scale 2 μm). **M)** Mitochondria total number. **N)** Mitochondria total perimeter. **O)** Mitochondria total size. **P)** Mitochondrial total aspect ratio **Q)** Mitochondrial total density from electron microscopy analyses, each point means one micrograph analyzed. Data presented as mean SEM. n=8-6 by group. A t-student test was performed. Statistical significance p=<0.05 *v/s CD. MCD: MCD: methionine-choline deficient diet; MCD+Ex: methionine-choline deficient diet + exercise.

To unveil the effect of exercise on NASH we analyzed LD-mitochondria interaction. MCD+Ex group decreased LD-mitochondria interaction compared to MCD (**Fig. 7A-D**) without changes in PLIN5 protein levels in the whole lysate (**Fig 6I-J**). Concerning changes in LD-M, the MCD+ex group decreased M-LD size compared to M-LD of the sedentary group (**Fig. 7E**). Interestingly, the MCD diet generated large M-LD (0.094 μm^2^ mean size MCD vs. 0.068 μm^2^ mean size HFD) that are closely related to PDM. Even though the MCD+exe group does not increase the elongated morphology of PDM compared to PDM from MCD mice, PDM shows markedly elongated morphology compared to CMs in both conditions (**Fig. 7F**). Finally, an absence of a correlation between LD-mitochondria interaction and collagen accumulation was observed (**Fig. 7G**).

**Figure 7.**
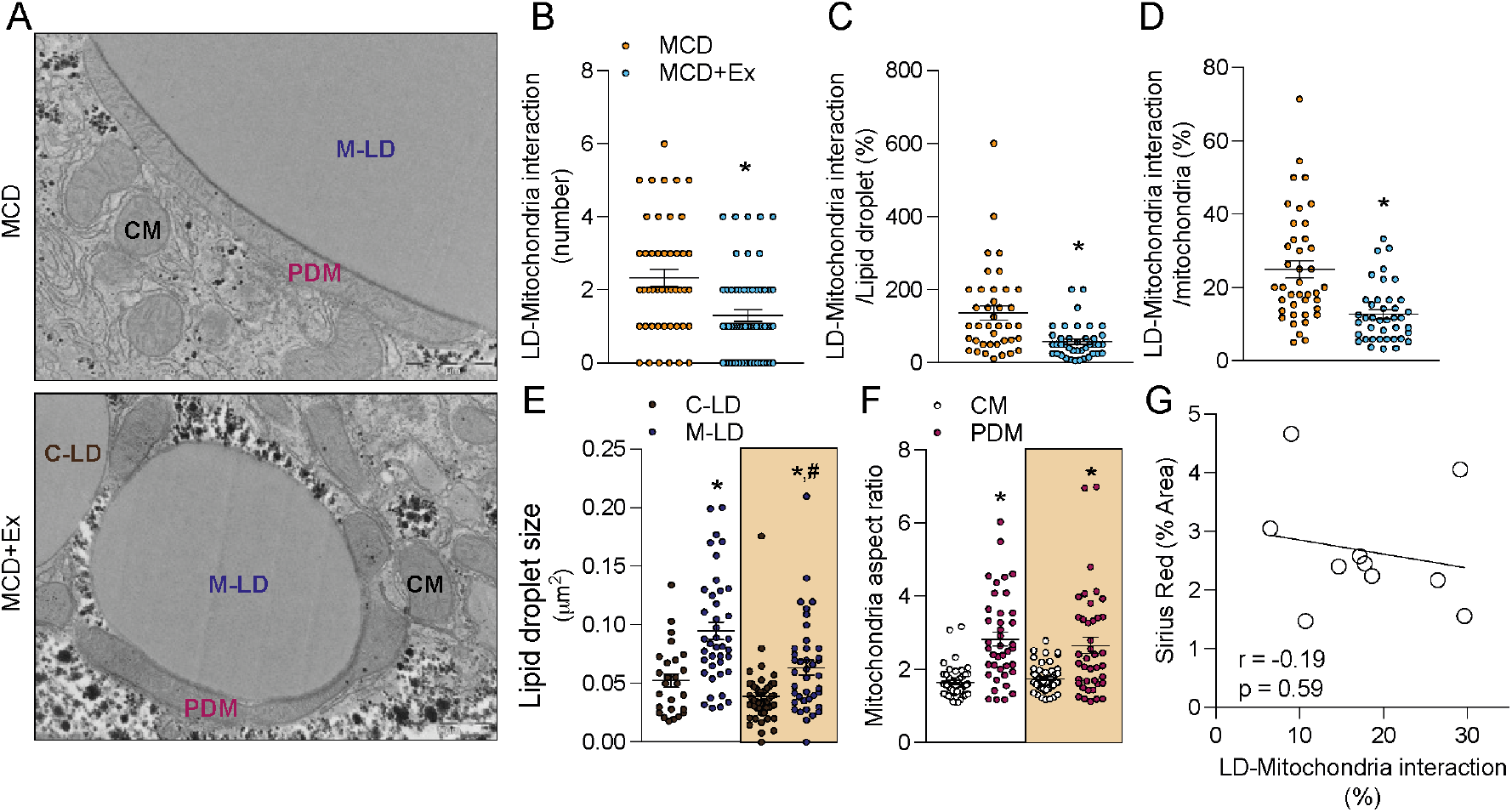
Exercise decreased Lipid droplet-mitochondria interaction in hepatocytes in MCD induced-NASH. **A)** Representative TEM micrograph of mice liver (25000x, bar scale 1 μm). **B)** Number of LD-mitochondria interactions. **C)** LD-Mitochondria interaction by the total number of mitochondria. **D)** LD-Mitochondria interaction by the total number of LD. **E)** LD size. **F)** Mitochondria aspect ratio. **G)** Correlation between LD-Mitochondria interaction/ total number of mitochondria (%) and Sirus Red. Data presented as mean SEM. n=8-6 by group. In A), C), D), E), F), each point means one micrograph analyzed. T-student test, 2-way ANOVA followed by Tukey*’*s post-test Pearson correlation coefficient were performed. Statistical significance p=<0.05; **\***vs. MCD or C-LD, ^**#**^vs. M-LD. MCD: MCD: methionine-choline deficient diet; MCD+Ex: methionine-choline deficient diet + exercise; CM: cytosolic mitochondria; PDM: peridroplet mitochondria; M-LD: Lipid droplets bound to mitochondria; C-LD: Lipid droplet non-bound to mitochondria.

## 4. Discussion

Despite several studies have shown that exercise attenuates NAFLD progression, the mechanism remains unclear. We showed that moderate treadmill exercise decreased LD-mitochondria interaction in hepatocytes and is associated with reduced NAFLD progression. Our results suggest that exercise decreases hepatic LD-mitochondria interaction, by diminishing the number of Peridroplet Mitochondria (PDM) and LD coupled to mitochondria (M-LD), which are associated with improvements in NAFLD.

Consistent with our results, studies show that exercise reduces NAFLD progression. In NAFL, 8 weeks of moderate treadmill exercise reduced liver weight, insulin resistance, serum triglycerides, and serum ALT transaminase in HFD-fed mice for 20 weeks (12). Likewise, 15 weeks of regular moderate treadmill exercise alleviated HFD-induced obesity, insulin intolerance, hyperlipidemia, and hyperglycemia (36). Similarly, 12 weeks of treadmill exercise reduced hepatic steatosis, NAFLD severity, ALT, and ASL transaminase levels and alleviated hepatic macrophage infiltration in mice fed for 24 weeks (37). In comparison to these studies, our study was a shorter training period (4 weeks v/s at least 8 weeks), proving the effectiveness of exercise in decreasing the progression of NAFLD even in a shorter time.

In addition, our results show that exercise reduces the severity of NASH. Consistently, 8 weeks of treadmill exercise (speed of 12.5 m/min for 60 min/5 times per week) attenuates the transition from fatty liver to steatohepatitis and reduces tumor formation in mice fed high-fat, choline-deficient diet (CD-HFD) for 20 weeks (4). Likewise, 3 weeks of treadmill exercise decreased hepatic steatosis in mice fed for 8 weeks on a low-fat, sucrose-rich, choline-deficient diet (5), and 12 weeks of treadmill exercise decreased collagen accumulation induced by the MCD diet (37).

Furthermore, exercise increases the frequency of small LD in both NAFL and NASH. This is relevant to NAFLD progression, where larger LD have been shown to be detrimental by increasing stellate cell activation, inflammation, and stress (33). In addition, it can be a beneficial adaptation to exercise. Small LD increases the contact surface area of LD, which can efficiently facilitate lipolysis by providing the lipases easy access to their lipid core (25). The small size of LD may result from high dynamics (a high rate of LD biogenesis and catabolism), as we described in exercised lean mice with normal liver (12). However, discordant studies have been described in NAFLD models, possibly due to increased LD biogenesis. Recently, a study showed that 15 weeks of moderate treadmill exercise in HFD-fed mice did not affect the expression of genes involved in LD biogenesis (e.g., dgat1, dgat2, or seipin) but did reduce the gene expression of genes involved in LD expansion (e.g., FITM2, CIDE A, and FSP27) (36). Therefore, exercise may induce a stop signal to LD expansion, resulting in small LD in NAFLD.

LD-mitochondria interaction was increased in our model of NAFL and NASH, both conditions of LD expansion. This observation is consistent with that reported in HFD-fed mice for 12 weeks (38) and contrasts with that reported in HFD-fed mice for 9 weeks in another study where no changes in LD-mitochondria interaction were observed (39). It is possible that LD-mitochondria interaction increases in relation to the severity of the NALFD. Thus, at 9 weeks of HFD is not different but at 12 weeks when the steatosis is higher, the LD-mitochondria interaction also increases. Moreover, studies in mice and humans with NAFLD show an increased abundance of PLIN5, a protein that promotes LD-mitochondria interaction (40).

Supporting the hypothesis that LD-mitochondria interaction promotes LD expansion in NAFL and NASH, we observed a positive correlation between LD-mitochondria interaction and mitochondria-bound LD size. This may also explain the positive correlation between LD-mitochondria interaction with ALT transaminase levels and the glucose tolerance test, as larger LDs are linked to worse NAFLD severity (17). In the HepG2 hepatocyte cell line, it has been described that oleic acid exposure increases LD-mitochondria interaction (21). Oleate exposure is known to generate large LD and LD expansion. In addition, PLIN5 overexpression in hepatocytes causes severe steatosis (20). In our models, the PDM show greater ATP synthesis capacity than CM, a feature associated with greater LD expansion in BAT (24). Finally, our model of MCD-induced NASH also increases LD-mitochondria interaction. Importantly, this model induces LD accumulation in the liver due to increased fatty acids uptake and inhibition of triglyceride export, without significantly affecting fatty acid oxidation (41). Furthermore, it generates larger mitochondria-bound LD (0.094 μm^2^ mean size MCD vs. 0.068 μm^2^ mean size HFD) and, in magnitude, more LD-mitochondria interactions (31.3% in MCD vs. 21.6% in HFD) than HFD. Taken together, these data suggest a role for LD-mitochondria interaction in LD expansion in NAFLD.

Whereas in white adipocytes, the LD-mitochondria-ER interaction promotes triglyceride synthesis from FA derived from *de novo* lipogenesis (42), in BAT the LD-mitochondria interaction promotes TAG synthesis, possibly, from esterification of available FA (24). Our data show that HFD induce-NAFL model promotes hepatic LD expansion from *de novo* lipogenesis and FA esterification in hepatocytes. However, our data from the MCD model could promotes the LD expansion by inhibiting FA export and increasing FA esterification without affecting *de novo* lipogenesis (41). Given that an increase in LD-mitochondria interaction was observed in this model, LD-mitochondria interaction may promote ATP synthesis from FA esterification in hepatocytes. Consistently, a hypothesis has been proposed in hepatocytes, in which LD-mitochondria interaction promotes FA esterification and ER-mitochondria interaction supports de novo lipogenesis and triglyceride export (35). It remains to be determined whether LD-mitochondria interaction promotes LD expansion by *de novo* lipogenesis, FA esterification, or both.

As a parallel with the skeletal muscle, exercise decreased contact among subsarcolemmal LD and mitochondria in diabetic, nonobese diabetic, and lean subjects (25). In this work our results showed that LD bound to mitochondria was related to a decreased size of subsarcolemmal LD. On the other hand, our results also show that under normal conditions, an increased LD-mitochondria interaction in the liver was observed under exercise. In this sense, skeletal muscle studies showed that endurance training increased LD-mitochondria interaction in healthy subjects (25,26,43). Therefore, it is possible that exercise has a differential effect on LD-mitochondria interaction, depending on health/disease status.

In addition, exercise modified each organelle involved in the interaction in our model. In addition, exercise decreased the size of LD bound to mitochondria, without changing the size of LD bound to non-mitochondria in NAFLD. This differential effect of exercise on LD subpopulations has been described in muscle. In this regard, exercise reduced the size of individual LDs located in the subsarcolemmal space in humans (25). Moreover, exercise reduced LDs in the subsarcolemmal region and increased LDs in the intermyofibrillar region of muscle in humans (43). Additionally, exercise increased the elongated morphology and ATP synthesis capacity of PDM. Considering that levels of mitochondrial complexes in PDMs were not changed, it is possible that exercise increases mitochondrial efficiency. In this regard, higher expression of supercomplex I+III has been reported in PDM versus CM in BAT (24). Exercise is known to induce the formation of mitochondrial supercomplexes and thus increase mitochondrial energy efficiency (44).

As we observed, exercise-induced changes in PDM also increased the ATP synthesis capacity of PDMs in NAFL. Consistently, a higher ATP synthesis capacity in PDM compared to CM was described in BAT (24). Theoretically, PDM generates ATP, which is used by acyl-CoA synthetases to bind fatty acid to coenzyme A (ATP-dependent reaction), which constitutes the first step of triglycerides synthesis and LD expansion (34). Therefore, exercise decreases LD-mitochondria interaction, decreasing PDM with lipogenic function, which induces a decrease in the expansion of LD bound to mitochondria in NAFLD. One hypothesis we cannot rule out is if exercise changes the function of LD-mitochondria interaction from LD expansion to LD utilization in NAFLD. In cells under starvation, the transfer of fatty acids from LD to mitochondria using a labeled fatty acid has been demonstrated (23,45). Furthermore, phosphorylation at serine 155 of PLIN5 in the liver induced by the protein kinase A pathway promotes LD-mitochondria interaction and enhances β-oxidation (46). Future studies will be required to determine whether the function of LD-mitochondria interaction can switch from LD expansion to LD utilization in the liver under different conditions.

This study has some limitations. First, it was not possible to isolate LD bound to mitochondria due to the strategy used allowed to isolate the total LD. Second, we did not measure other anchoring proteins described to also participate in the LD-mitochondria interaction (e.g., DGAT2, MIGA-2). However, we directly measured the physical contact between both organelles by electron microscopy. Third, we did not perform other types of exercise training (e.g., resistance training). Therefore, this work*’*s conclusions could only be related to aerobic treadmill exercise. Finally, we only included male mice in this study, which limits the generalizability of the results to women.

## 5. Conclusion

Here, we showed that aerobic exercise decreases LD-mitochondria interaction in hepatocytes and is associated with reduced progression of NAFLD. We propose that exercise changes hepatic LD-mitochondria interaction, improving NAFLD.

## Contribution statement

JCB, MIHA, VC, and RT conceived the study. JCB, FD-C, FP, KE, AMF, IM-R, VH, IL-S, and RV performed the experiments. JCB, RT, and VC analyzed the data. JCB and RT wrote the manuscript and all authors contributed to editing.

## Acknowledgments

We thank Alba Fajardo from Cell Compartments and Signaling Group, Institut d’Investigacions Biomèdiques August Pi i Sunyer (IDIBAPS), Barcelona, Spain; Microscopy Facility Universidad Católica; Chile; CCIT of Universitat de Barcelona; Histology Unit, ACCDIS, University of Chile; Izan Cabrerizo from Universitat de Barcelona.

## Supplementary figures

**Supplementary figure 1.**
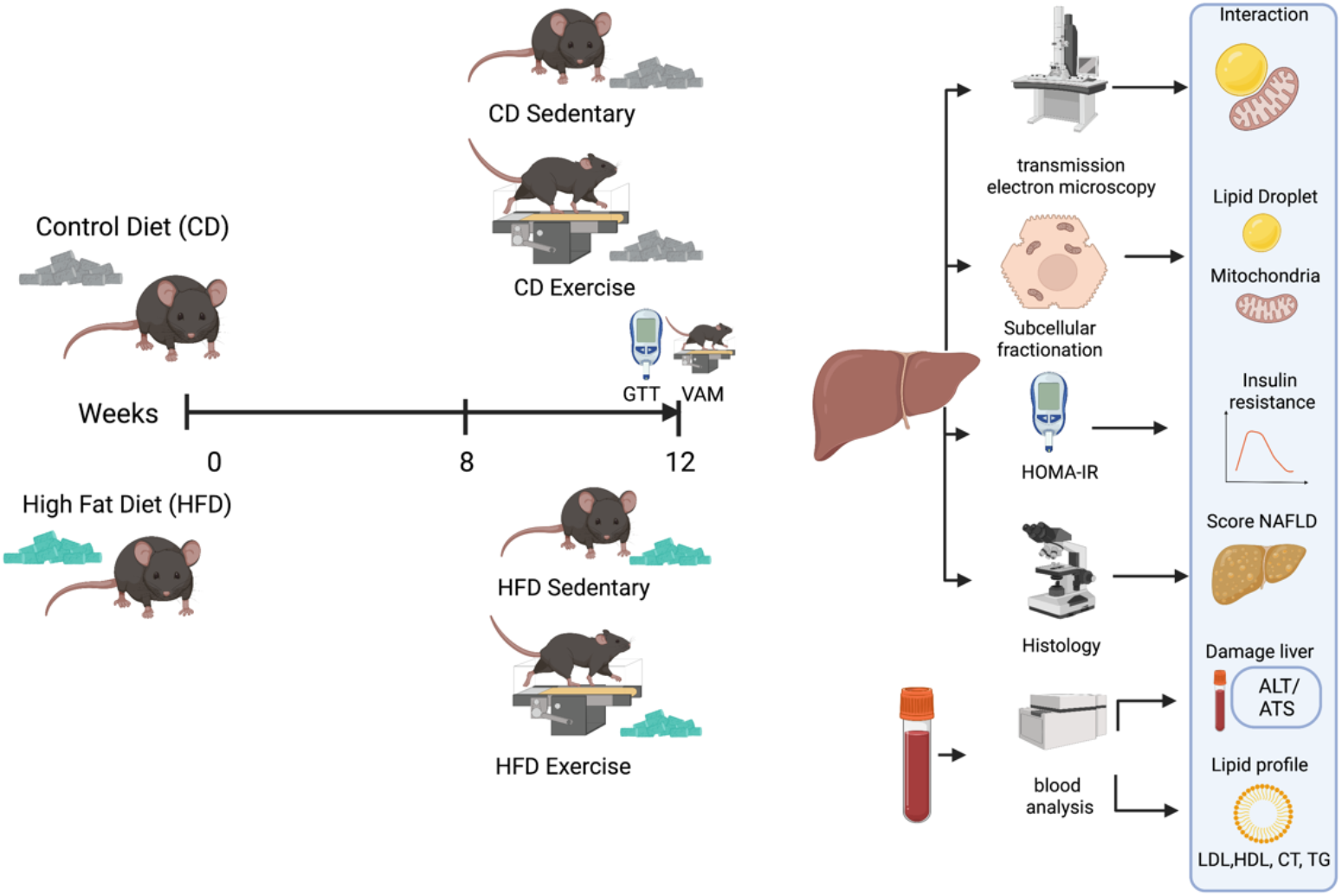
Cohort 1 study design.

**Supplementary figure 2.**
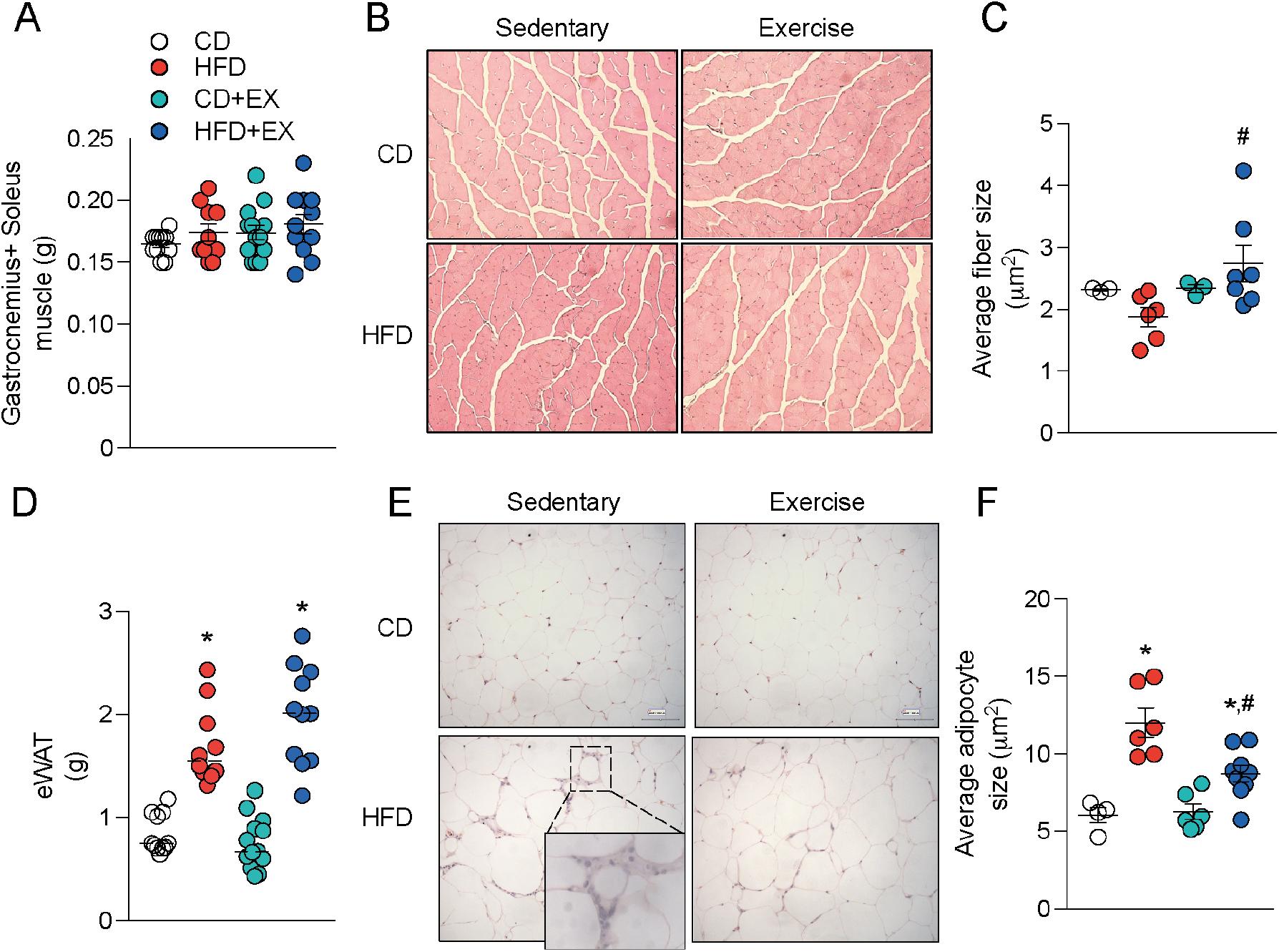
Exercise increased muscle fiber size and reduced adipocyte size in HFD-induced NAFLD. A) Gastronemius and Soleus muscle weight. B) Skeletal muscle (H&E stain histology, 20x). C) Average fiber size (Gastronemius). D) epididymal white adipose tissue weight (eWAT). E) eWAT (H&E stain histology, 20x). F) Average adipocyte size (eWAT). Data presented as mean SEM. n=9-13 mice per group. 2-way ANOVA followed by Tukey*’*s post-test. Statistical significance p=<0.05; *vs CD, ^&^vs CD, #vs HFD. CD: control diet; CD+Ex: control diet + exercise; HFD: high-fat diet; HFD+Ex: high-fat diet + exercise.

**Supplementary figure 3.**
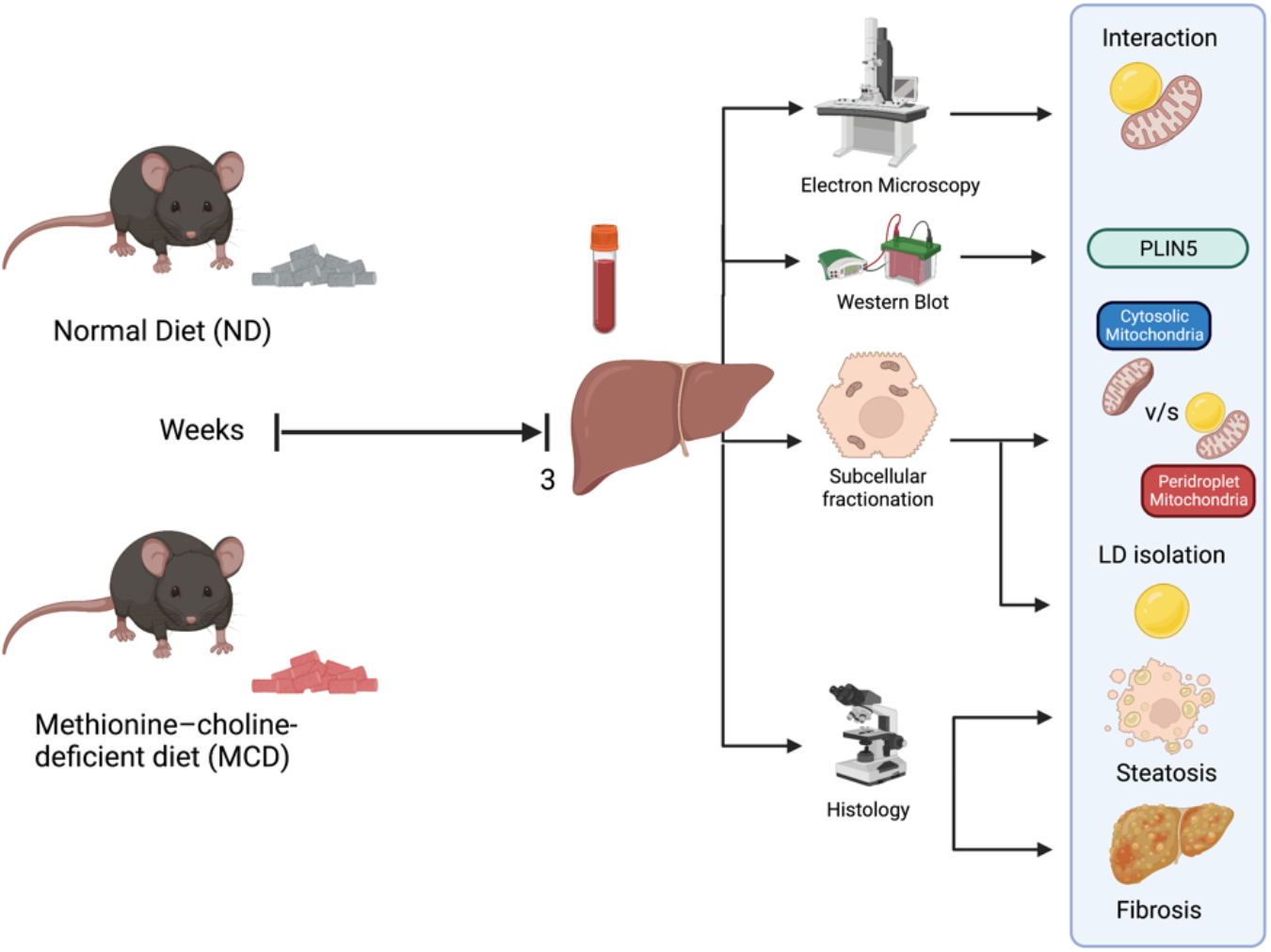
Cohort 2 study design.

**Supplementary figure 4.**
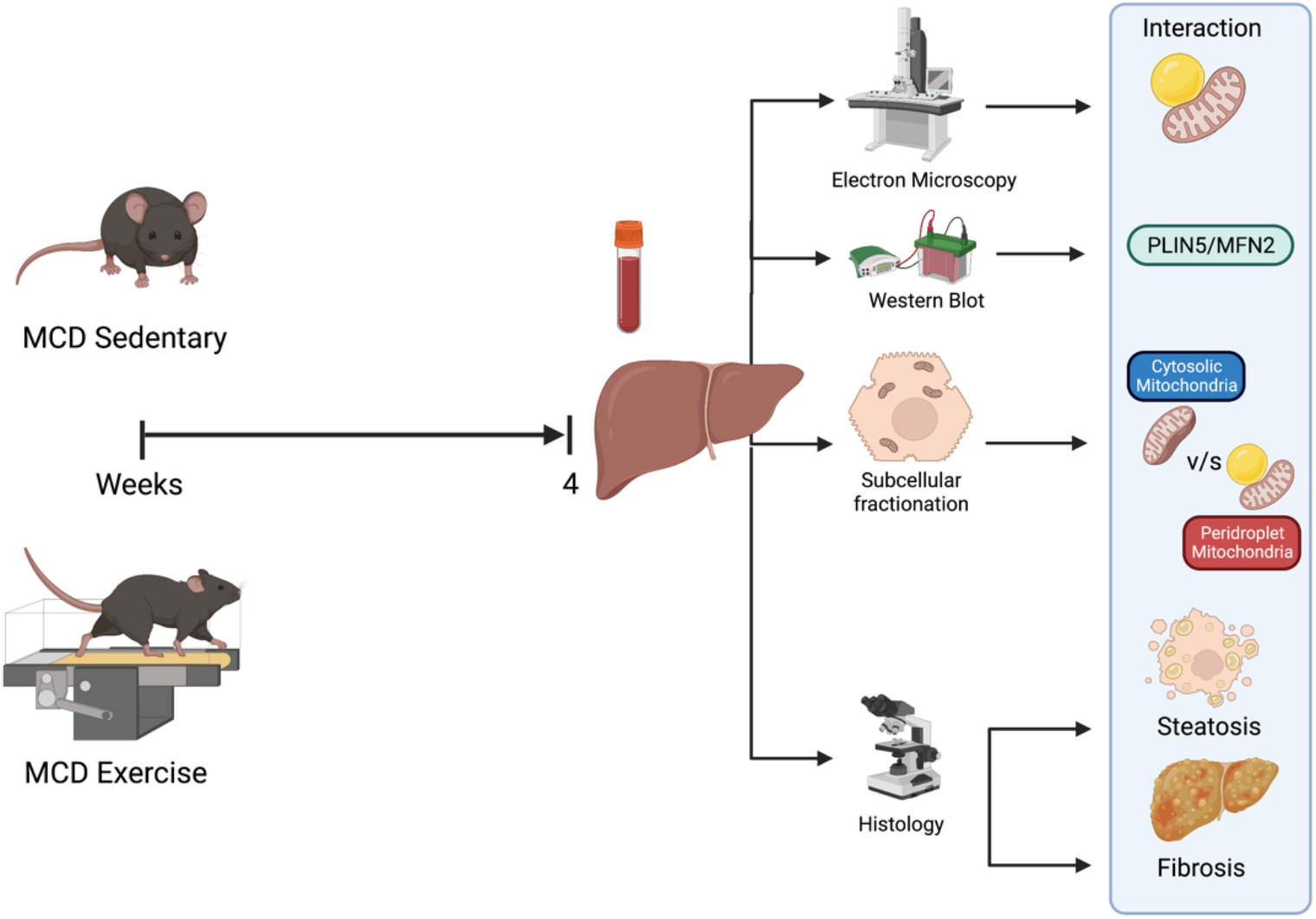
Cohort 3 study design.

## References

1. Riazi K, Azhari H, Charette JH, Underwood FE, King JA, Afshar EE, Swain MG, Congly SE, Kaplan GG, Shaheen A-A. The prevalence and incidence of NAFLD worldwide: a systematic review and meta-analysis. Lancet Gastroenterol Hepatol (2022) 7:851–861. doi: 10.1016/S2468-1253(22)00165-0

2. Targher G, Byrne CD, Tilg H. NAFLD and increased risk of cardiovascular disease: clinical associations, pathophysiological mechanisms and pharmacological implications. Gut (2020) 69:1691–1705. doi: 10.1136/gutjnl-2020-320622

3. Human Fatty Liver Disease: Old Questions and New Insights | Science. https://www.science.org/doi/10.1126/science.1204265?url_ver=Z39.88-2003&rfr_id=ori:rid:crossref.org&rfr_dat=cr_pub%20%200pubmed [Accessed January 14, 2023]

4. Guarino M, Kumar P, Felser A, Terracciano LM, Guixé-Muntet S, Humar B, Foti M, Nuoffer J-M, St-Pierre MV, Dufour J-F. Exercise Attenuates the Transition from Fatty Liver to Steatohepatitis and Reduces Tumor Formation in Mice. Cancers (2020) 12:1407. doi: 10.3390/cancers12061407

5. Alex S, Boss A, Heerschap A, Kersten S. Exercise training improves liver steatosis in mice. Nutr Metab (2015) 12:29. doi: 10.1186/s12986-015-0026-1

6. Fredrickson G, Barrow F, Dietsche K, Parthiban P, Khan S, Robert S, Demirchian M, Rhoades H, Wang H, Adeyi O, et al. Exercise of high intensity ameliorates hepatic inflammation and the progression of NASH. Mol Metab (2021) 53:101270. doi: 10.1016/j.molmet.2021.101270

7. Linden MA, Sheldon RD, Meers GM, Ortinau LC, Morris EM, Booth FW, Kanaley JA, Vieira-Potter VJ, Sowers JR, Ibdah JA, et al. Aerobic exercise training in the treatment of non-alcoholic fatty liver disease related fibrosis. J Physiol (2016) 594:5271– 5284. doi: 10.1113/JP272235

8. Pino-de la Fuente F, Bórquez JC, Díaz-Castro F, Espinosa A, Chiong M, Troncoso R. Exercise regulation of hepatic lipid droplet metabolism. Life Sci (2022) 298:120522. doi: 10.1016/j.lfs.2022.120522

9. Brouwers B, Hesselink MKC, Schrauwen P, Schrauwen-Hinderling VB. Effects of exercise training on intrahepatic lipid content in humans. Diabetologia (2016) 59:2068– 2079. doi: 10.1007/s00125-016-4037-x

10. Brouwers B, Schrauwen-Hinderling VB, Jelenik T, Gemmink A, Sparks LM, Havekes B, Bruls Y, Dahlmans D, Roden M, Hesselink MKC, et al. Exercise training reduces intrahepatic lipid content in people with and people without nonalcoholic fatty liver. Am J Physiol Endocrinol Metab (2018) 314:E165–E173. doi: 10.1152/ajpendo.00266.2017

11. Marcinko K, Sikkema SR, Samaan MC, Kemp BE, Fullerton MD, Steinberg GR. High intensity interval training improves liver and adipose tissue insulin sensitivity. Mol Metab (2015) 4:903–915. doi: 10.1016/j.molmet.2015.09.006

12. la Fuente FP, Quezada L, Sepúlveda C, Monsalves-Alvarez M, Rodríguez JM, Sacristán C, Chiong M, Llanos M, Espinosa A, Troncoso R. Exercise regulates lipid droplet dynamics in normal and fatty liver. Biochim Biophys Acta BBA - Mol Cell Biol Lipids (2019) 1864:158519. doi: 10.1016/j.bbalip.2019.158519

13. Winn NC, Liu Y, Rector RS, Parks EJ, Ibdah JA, Kanaley JA. Energy-matched moderate and high intensity exercise training improves nonalcoholic fatty liver disease risk independent of changes in body mass or abdominal adiposity - A randomized trial. Metabolism (2018) 78:128–140. doi: 10.1016/j.metabol.2017.08.012

14. Ok D-P, Ko K, Bae JY. Exercise without dietary changes alleviates nonalcoholic fatty liver disease without weight loss benefits. Lipids Health Dis (2018) 17:207. doi: 10.1186/s12944-018-0852-z

15. Gluchowski NL, Becuwe M, Walther TC, Farese RV. Lipid droplets and liver disease: from basic biology to clinical implications. Nat Rev Gastroenterol Hepatol (2017) 14:343–355. doi: 10.1038/nrgastro.2017.32

16. Olzmann JA, Carvalho P. Dynamics and functions of lipid droplets. Nat Rev Mol Cell Biol (2019) 20:137–155. doi: 10.1038/s41580-018-0085-z

17. Mashek DG. Hepatic lipid droplets: A balancing act between energy storage and metabolic dysfunction in NAFLD. Mol Metab (2021) 50:101115. doi: 10.1016/j.molmet.2020.101115

18. Valm AM, Cohen S, Legant WR, Melunis J, Hershberg U, Wait E, Cohen AR, Davidson MW, Betzig E, Lippincott-Schwartz J. Applying systems-level spectral imaging and analysis to reveal the organelle interactome. Nature (2017) 546:162–167. doi: 10.1038/nature22369

19. Keenan SN, Meex RC, Lo JCY, Ryan A, Nie S, Montgomery MK, Watt MJ. Perilipin 5 Deletion in Hepatocytes Remodels Lipid Metabolism and Causes Hepatic Insulin Resistance in Mice. Diabetes (2019) 68:543–555. doi: 10.2337/db18-0670

20. Trevino MB, Mazur-Hart D, Machida Y, King T, Nadler J, Galkina EV, Poddar A, Dutta S, Imai Y. Liver Perilipin 5 Expression Worsens Hepatosteatosis But Not Insulin Resistance in High Fat-Fed Mice. Mol Endocrinol Baltim Md (2015) 29:1414–1425. doi: 10.1210/me.2015-1069

21. Eynaudi A, Díaz-Castro F, Bórquez JC, Bravo-Sagua R, Parra V, Troncoso R. Differential Effects of Oleic and Palmitic Acids on Lipid Droplet-Mitochondria Interaction in the Hepatic Cell Line HepG2. Front Nutr (2021) 8: https://www.frontiersin.org/articles/10.3389/fnut.2021.775382 [Accessed January 25, 2023]

22. Wang H, Sreenivasan U, Hu H, Saladino A, Polster BM, Lund LM, Gong D-W, Stanley WC, Sztalryd C. Perilipin 5, a lipid droplet-associated protein, provides physical and metabolic linkage to mitochondria. J Lipid Res (2011) 52:2159–2168. doi: 10.1194/jlr.M017939

23. Miner GE, So CM, Edwards W, Herring LE, Coleman RA, Klett EL, Cohen S. Perilipin 5 interacts with Fatp4 at membrane contact sites to promote lipid droplet-to-mitochondria fatty acid transport. (2022)2022.02.03.479028. doi: 10.1101/2022.02.03.479028

24. Benador IY, Veliova M, Mahdaviani K, Petcherski A, Wikstrom JD, Assali EA, Acín-Pérez R, Shum M, Oliveira MF, Cinti S, et al. Mitochondria Bound to Lipid Droplets Have Unique Bioenergetics, Composition, and Dynamics that Support Lipid Droplet Expansion. Cell Metab (2018) 27:869-885.e6. doi: 10.1016/j.cmet.2018.03.003

25. de Almeida ME, Nielsen J, Petersen MH, Wentorf EK, Pedersen NB, Jensen K, Højlund K, Ørtenblad N. Altered intramuscular network of lipid droplets and mitochondria in type 2 diabetes. Am J Physiol Cell Physiol (2023) 324:C39–C57. doi: 10.1152/ajpcell.00470.2022

26. Tarnopolsky MA, Rennie CD, Robertshaw HA, Fedak-Tarnopolsky SN, Devries MC, Hamadeh MJ. Influence of endurance exercise training and sex on intramyocellular lipid and mitochondrial ultrastructure, substrate use, and mitochondrial enzyme activity. Am J Physiol Regul Integr Comp Physiol (2007) 292:R1271–1278. doi: 10.1152/ajpregu.00472.2006

27. Bankhead P, Loughrey MB, Fernández JA, Dombrowski Y, McArt DG, Dunne PD, McQuaid S, Gray RT, Murray LJ, Coleman HG, et al. QuPath: Open source software for digital pathology image analysis. Sci Rep (2017) 7:16878. doi: 10.1038/s41598-017-17204-5

28. Hernández-Alvarez MI, Sebastián D, Vives S, Ivanova S, Bartoccioni P, Kakimoto P, Plana N, Veiga SR, Hernández V, Vasconcelos N, et al. Deficient Endoplasmic Reticulum-Mitochondrial Phosphatidylserine Transfer Causes Liver Disease. Cell (2019) 177:881-895.e17. doi: 10.1016/j.cell.2019.04.010

29. Tapia PJ, Figueroa A-M, Eisner V, González-Hódar L, Robledo F, Agarwal AK, Garg A, Cortés V. Absence of AGPAT2 impairs brown adipogenesis, increases IFN stimulated gene expression and alters mitochondrial morphology. Metabolism (2020) 111:154341. doi: 10.1016/j.metabol.2020.154341

30. Acín-Perez R, Petcherski A, Veliova M, Benador IY, Assali EA, Colleluori G, Cinti S, Brownstein AJ, Baghdasarian S, Livhits MJ, et al. Recruitment and remodeling of peridroplet mitochondria in human adipose tissue. Redox Biol (2021) 46:102087. doi: 10.1016/j.redox.2021.102087

31. Mammalian lipid droplets are innate immune hubs integrating cell metabolism and host defense | Science. https://www.science.org/doi/10.1126/science.aay8085?url_ver=Z39.88-2003&rfr_id=ori:rid:crossref.org&rfr_dat=cr_pub%20%200pubmed [Accessed January 15, 2023]

32. Ngo J, Benador IY, Brownstein AJ, Vergnes L, Veliova M, Shum M, Acín-Pérez R, Reue K, Shirihai OS, Liesa M. Isolation and functional analysis of peridroplet mitochondria from murine brown adipose tissue. STAR Protoc (2021) 2:100243. doi: 10.1016/j.xpro.2020.100243

33. Scorletti E, Carr RM. A new perspective on NAFLD: Focusing on lipid droplets. J Hepatol (2021)S0168827821021814. doi: 10.1016/j.jhep.2021.11.009

34. Benador IY, Veliova M, Liesa M, Shirihai OS. Mitochondria Bound to Lipid Droplets: Where Mitochondrial Dynamics Regulate Lipid Storage and Utilization. Cell Metab (2019) 29:827–835. doi: 10.1016/j.cmet.2019.02.011

35. Shum M, Ngo J, Shirihai OS, Liesa M. Mitochondrial oxidative function in NAFLD: Friend or foe? Mol Metab (2021) 50:101134. doi: 10.1016/j.molmet.2020.101134

36. Yang Y, Li X, Liu Z, Ruan X, Wang H, Zhang Q, Cao L, Song L, Chen Y, Sun Y. Moderate Treadmill Exercise Alleviates NAFLD by Regulating the Biogenesis and Autophagy of Lipid Droplet. Nutrients (2022) 14:4910. doi: 10.3390/nu14224910

37. Exercise suppresses NLRP3 inflammasome activation in mice with diet-induced NASH: a plausible role of adropin | Elsevier Enhanced Reader. doi: 10.1038/s41374-020-00508-y

38. Krahmer N, Najafi B, Schueder F, Quagliarini F, Steger M, Seitz S, Kasper R, Salinas F, Cox J, Uhlenhaut NH, et al. Organellar Proteomics and Phospho-Proteomics Reveal Subcellular Reorganization in Diet-Induced Hepatic Steatosis. Dev Cell (2018) 47:205-221.e7. doi: 10.1016/j.devcel.2018.09.017

39. Khan SA, Wollaston-Hayden EE, Markowski TW, Higgins L, Mashek DG. Quantitative analysis of the murine lipid droplet-associated proteome during diet-induced hepatic steatosis. J Lipid Res (2015) 56:2260–2272. doi: 10.1194/jlr.M056812

40. Wang C, Zhao Y, Gao X, Li L, Yuan Y, Liu F, Zhang L, Wu J, Hu P, Zhang X, et al. Perilipin 5 improves hepatic lipotoxicity by inhibiting lipolysis. Hepatol Baltim Md (2015) 61:870–882. doi: 10.1002/hep.27409

41. Rinella ME, Elias MS, Smolak RR, Fu T, Borensztajn J, Green RM. Mechanisms of hepatic steatosis in mice fed a lipogenic methionine choline-deficient diet. J Lipid Res (2008) 49:1068–1076. doi: 10.1194/jlr.M800042-JLR200

42. Freyre CAC, Rauher PC, Ejsing CS, Klemm RW. MIGA2 Links Mitochondria, the ER, and Lipid Droplets and Promotes De Novo Lipogenesis in Adipocytes. Mol Cell (2019) 76:811-825.e14. doi: 10.1016/j.molcel.2019.09.011

43. Devries MC, Samjoo IA, Hamadeh MJ, McCready C, Raha S, Watt MJ, Steinberg GR, Tarnopolsky MA. Endurance training modulates intramyocellular lipid compartmentalization and morphology in skeletal muscle of lean and obese women. J Clin Endocrinol Metab (2013) 98:4852–4862. doi: 10.1210/jc.2013-2044

44. Casuso RA, Al-Fazazi S, Hidalgo-Gutierrez A, López LC, Plaza-Díaz J, Rueda-Robles A, Huertas JR. Hydroxytyrosol influences exercise-induced mitochondrial respiratory complex assembly into supercomplexes in rats. Free Radic Biol Med (2019) 134:304–310. doi: 10.1016/j.freeradbiomed.2019.01.027

45. Rambold AS, Cohen S, Lippincott-Schwartz J. Fatty acid trafficking in starved cells: regulation by lipid droplet lipolysis, autophagy, and mitochondrial fusion dynamics. Dev Cell (2015) 32:678–692. doi: 10.1016/j.devcel.2015.01.029

46. Keenan SN, De Nardo W, Lou J, Schittenhelm RB, Montgomery MK, Granneman JG, Hinde E, Watt MJ. Perilipin 5 S155 phosphorylation by PKA is required for the control of hepatic lipid metabolism and glycemic control. J Lipid Res (2021) 62:100016. doi: 10.1194/jlr.RA120001126

